# Mismatch Negativity Predicts Pattern Separation

**DOI:** 10.1101/2020.05.19.102707

**Authors:** Deena Herman, Stevenson Baker, Jaime Cazes, Claude Alain, R. Shayna Rosenbaum

**Author notes:** Contact: Stevenson Baker, Department of Psychology, Faculty of Health, York University, 4700 Keele Street, Toronto, ON M3J 1P3, CANADA. (or) R. Shayna Rosenbaum, Department of Psychology, Faculty of Health, York University, 4700 Keele Street, Toronto, ON M3J 1P3, CANADA.

## Abstract

We are so tuned to sensory changes that we can detect novelty within hundreds of milliseconds. To what extent does our capacity to automatically discriminate auditory inputs influence encoding of long-term memories? We recorded mismatch negativity (MMN), an event-related potential (ERP) indexing perceptual discrimination, as participants were presented with sound patterns while watching a muted movie. MMN strength predicted how well naïve listeners separated the previously heard from new micropatterns on a subsequent recognition test, providing evidence that the MMN translates into mnemonic pattern separation. Our investigation is the first to show that our capacity to discriminate auditory inputs, as measured by MMN, gives rise to unique memories.

---

It has been over 40 years since a Finnish research team discovered that our brain’s ability to discriminate auditory stimuli can be indexed by the mismatch negativity (MMN) signature (1). It reveals itself in EEG experiments whenever an oddball event violates predictions established from the preceding stimuli. These auditory oddballs generate an MMN response 100–200 ms after the onset of the deviant stimulus (2). Over the decades, investigators have found the MMN to be a reliable change-detection event-related potential (ERP) component (3). Researchers are now beginning to establish that the MMN response might transcend its acknowledged role as an index of perceptual discrimination and relate to higher-order cognitive processes, such as long-term memory (3). We propose that one way the MMN does so is through interacting with the neurobiological mechanism of pattern separation, a memory process by which similar or overlapping information is disambiguated into unique events at encoding (4).

There are hints in the literature that the MMN interacts with pattern separation or with pattern completion (recollection of memories from partial cues) (4). For example, data from animal models suggest that mismatch responses manifested within the auditory cortex are detected by the hippocampus (5), an area essential for episodic memory (6) possibly within the dentate gyrus (DG) or CA1/CA3. These hippocampal subfields have been implicated in the sparse coding on which pattern separation depends (2, 7). Additional links can be inferred from human studies showing that the neuronal mechanisms responsible for MMN generation may be explained by a model of hierarchical inference (i.e., predictive coding and predictive error) (2, 7). Indeed, a leading theory of the MMN response is that it involves the interplay of predictive coding — or top-down perceptual inferences — with prediction error, or neural responses following violations of expected inputs (2), including in the hippocampus (8, 9). Furthermore, hippocampal indices of predictive coding/predictive error are being investigated in contemporary studies of behavioral pattern separation (also known as mnemonic discrimination) and pattern completion (10, 11).

In humans, behavioral pattern separation is classically illustrated by assessing participants’ ability to differentiate studied from unstudied objects (12). Unclear is the extent to which mnemonic discrimination applies to the auditory domain or to unknown abstract stimuli. Experiments used to investigate the MMN offer an incidental learning situation involving unique patterns of stimuli that vary in a systematic, quantifiable way. We predicted that the brain dynamics following mismatch prediction errors would facilitate recognition memory and behavioral pattern separation of abstract sounds, thereby serving a broader purpose beyond perception.

To determine if the MMN relates to behavioral discrimination, healthy adults (*n*=31, 18–32 years, 18 women) participated in an experiment that included passive listening at study and a subsequent recognition task. The stimuli used to generate the MMN were 500 ms micropatterns (13), constructed using a sequence of five 100 ms tones. During the passive listening phase, we presented one micropattern 70% of the time (700 trials). It was designated the “standard.” A second micro-pattern, delivered 30% of the time (300 trials), was the de facto “deviant.” We played the micropatterns while participants watched 25 minutes of a muted movie (*Toy Story*) (14).

We anticipated that the presentation of the unexpected deviant would elicit a negative deflection in the ERP waveform (MMN), calculated as the difference between the standard and deviant waveforms (3). We found the MMN waveform was largest at six frontocentral electrodes with two peaks, MMN1 and MMN2. Using an iterative application of low-resolution electromagnetic tomography (BESA v7.0) we found standard and deviant sounds were associated with source activity in the auditory cortex of the superior temporal gyrus. The contrast in activity between the standard and deviant micropatterns, however, revealed greater source activity for the deviant in the superior temporal gyrus and the medial temporal lobe (Figure 2C).

**Figure 1.**
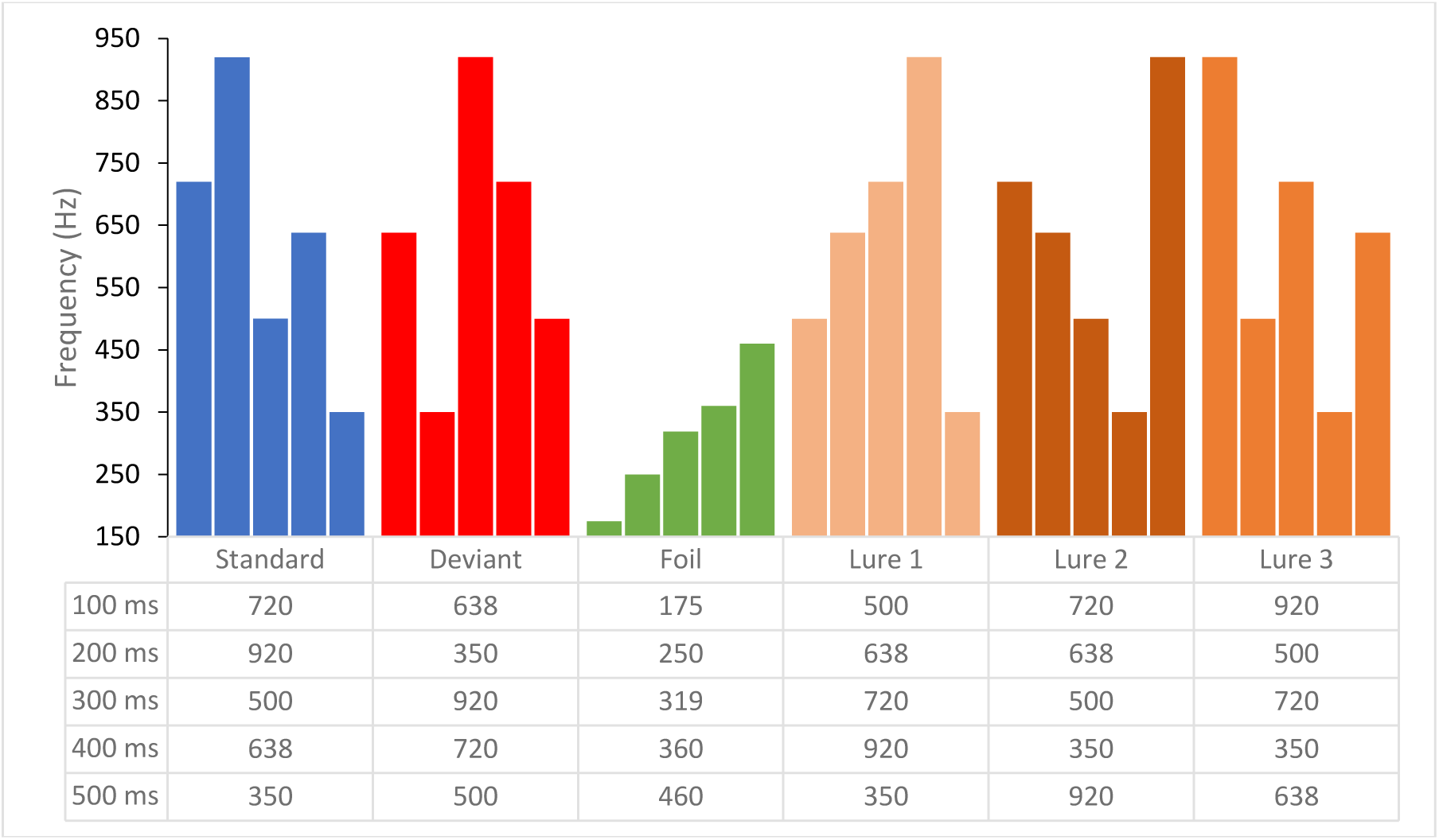
Schematic illustration of the 500-ms standard, deviant, foil and lure micropatterns.

**Figure 2.**
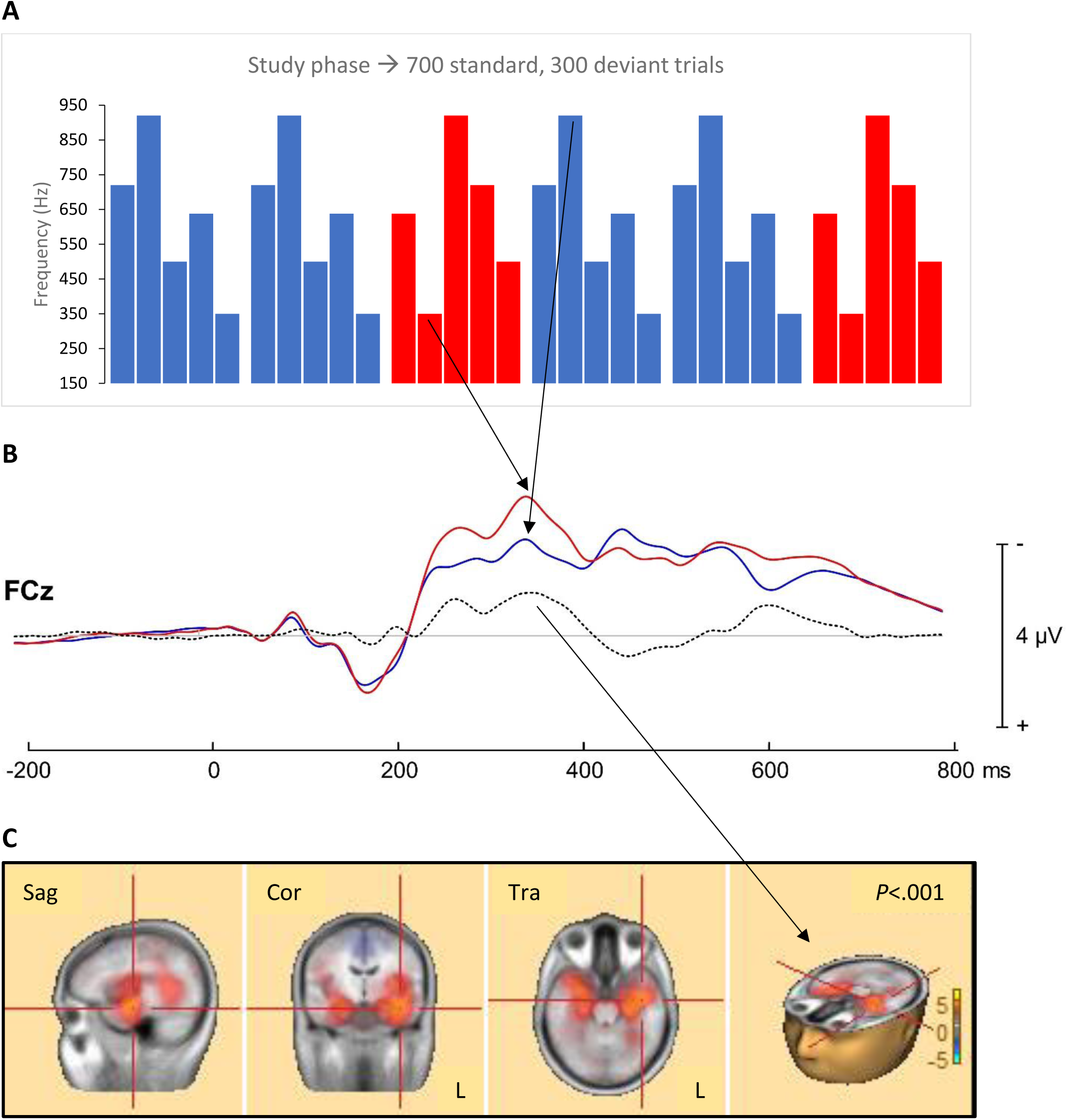
Micropatterns at Study. Notes: (A) Schematic illustration of the 500-ms standard and deviant micropatterns. (B) Grand average frontal-central pole (FCz) event-related potentials in response to the standard (blue line) and the deviant (red line) micropatterns. Black arrows indicate the 200 ms tone; detection of the mismatch here is responsible for the MMN1 amplitude spike at approximately 350 ms, which is 150 ms after the onset of the mismatch with the deviant micropattern. The dotted black line indicates the MMN waveform. (C) Four views of the CLARA source analysis of the difference in activity between deviant and standard conditions (300-400 ms latency). Red and yellow shading represents areas of the brain at this latency where deviant micropattern activity is significantly greater (*p*<.001) than standard micropattern activity. Sag, sagittal; Cor, coronal; Tra, transverse; L, right.

During the test phase, participants were presented with the standard and deviant (old) micropatterns from the passive listening phase, randomly intermixed with four unstudied (new) micropatterns. Three of the new micropatterns were similar to the standard and deviant and were designated as lures. We labeled a fourth micropattern — more distinct in pitch and temporal arrangement — as the foil. The test phase consisted of 60 trials (6 micropatterns x 10 trials each), but, as we could not find a significant difference in recognition memory for the three lures, they were treated as a single condition. For each trial, participants were instructed to indicate with a button press whether they had heard the micropattern while watching *Toy Story* (14).

Recognition accuracy for old (combined standard and deviant) micropatterns was above chance. On average, participants could correctly identify 91.61% of the standard and deviant micropatterns as old, and showed additional high accuracy for foils, correctly identifying 93.55% as new. Thus, we demonstrated the ability of naïve listeners to discriminate unique acoustic patterns in memory.

Correct rejection rates for the lures (*M* = 58.84%; *SE* = 3.98%), however, were significantly lower than they were for foils (*M* = 93.55%, *SE* = 2.61%). This difference, –34.71%, 95% CI [–.43.34%, –.26.08%], was significant, two-tailed *t*-test, *t*(30) = –8.22, *p* <.001, reflecting the greater interference involved in the lure condition. We likewise calculated *d’* for old relative to new micropatterns. We found that participants were significantly more accurate at recognizing old micropatterns relative to the foil, *d’* (O,F), *M* = 2.88, *SE* = .13, than they were at identifying old micropatterns relative to the lures, *d’* (O,L), *M* = 1.74, *SE* = .14. This difference, 1.15, 95% CI [.87, 1.42], was significant, two-tailed *t*-test, *t*(30) = 8.51, *p* < .001, *r* = .84.

To disentangle the MMN representation of behavioral pattern separation, we correlated the first and highest MMN peak (MMN1) amplitudes with *d’* scores. We found the MMN1 was significantly related to lure discrimination, *d’* (O,L), *r* = –.40, 95% BCa CI [–.69, –.02], *p* = .013, one-tailed. After correcting for multiple comparisons, we failed to find a significant relationship between the MMN1 and foil recognition, *d’* (O,F), *r* = –.33, 95% BCa CI [–.55, –.07], *p* = .036, one-tailed. This primary finding that the MMN1 and *d’* (O,L) are correlated is consistent with our prediction that strong mnemonic representation is formed from the MMN and that it predicts behavioral pattern separation.

We propose that the MMN amplitude generated by the abstract micropatterns may index an interaction of predictive coding within the auditory cortex and prediction errors within the hippocampus. A precedent for this assertion may be found in vision science. Here, prediction errors in response to violations of perceptual expectations have been found to bias hippocampal CA3 and CA1 subfields (10). These biases lead to “states,” whereby the hippocampus is more conducive to pattern separation or pattern completion (10). In our study, predictive coding of auditory inputs, encouraged by CA3 pattern completion, would presumably help to decode memories of old micropatterns, and predictive errors would bias the CA1 towards sparse encoding of highly similar lures (9, 10).

Results from our source analysis suggest that the auditory cortex and medial temporal lobes contribute to the MMN signal. Ruusuvirta (5) showed that the MMN signal in rodents is elicited in both the auditory cortex and the hippocampal DG, subiculum, and CA1. The CA1 receives input from the DG via the CA3 (4). It also back projects onto the neocortex to activate modality-specific cortical areas involved in episodic memory (4, 11). Thus, the auditory cortical source of the MMN might represent the back projections from the CA1 to the auditory cortex in service of the representation of the deviant in memory (15), in the same way abstract visual information is hypothesized to interact with the visual cortex (11).

Our findings of a significant correlation between the MMN amplitude for pre-experimentally unknown, incidentally encoded auditory stimuli and behavioral discrimination of micropatterns support our position that the MMN is a direct measure of auditory pattern separation at encoding. To our knowledge, this is the first study to examine the nature of pattern separation with novel auditory stimuli and to describe the MMN as a neural signature of mnemonic discrimination. A leading view is that the MMN is an overall barometer of neuronal dynamics and brain plasticity (2, 3). Our study suggests that such perceptual and mnemonic malleability is a manifestation of pattern separation, predictive coding, and match-mismatch detection — all different iterations of similar neurophysiological, hippocampally dependent processes.

## Methods

Methods and additional references are available in the Supplementary Information.

## Acknowledgments

The authors thank Yasha Amani, Ricky Chow, Jad Daou, and Yarden Levy for their help and support in this project. This study was funded by the Canada First Research Excellence Fund and York Research Chair to R.S.R.

## Author contributions

S.B., C.A., and R.S.R. conceived the project, and D.H., S.B., J.C., C.A. and R.S.R. designed the experiment. Data collection and analysis were performed by D.H., S.B., C.A., and R.S.R. The manuscript was drafted by D.H., S.B., C.A. and R.S.R. All authors reviewed and approved the final draft.

## Data availability

The data that support the findings of this study are available on request from the corresponding author (R.S.R.).

**Correspondence and requests for materials** should be addressed to R.S.R. or S.B.

## Supplementary Information

### Mismatch Negativity Predicts Pattern Separation

#### Participants

Thirty-six individuals (19 females and 17 males, aged 18 to 32 years) participated in the study. We excluded five participants from final data analysis for the following reasons: one fell asleep during EEG recording, one had pitch discrimination deficits, one was left-handed, one had a shorter interstimulus interval than other participants that could have affected performance, and one failed to understand or disregarded test instructions. Data analysis, therefore, included 31 individuals (18 females and 13 males, aged 18 to 32 years). This sample size is similar to previous investigations using a comparable age range (11, 16, 17). All participants were right-handed, and none reported learning disabilities or neuropsychological disorders. Participants were screened for anxiety and depression symptoms using the Generalized Anxiety Disorder 7-item scale (GAD7) and the Patient Health Questionnaire Depression 8-item scale (PHQ8). Three participants reported a current diagnosis of a mood disorder, but only one of the three exceeded the severity threshold of 15 on both tests. Before ERP acquisition, we administered an audiogram; all participants had normal pure-tone thresholds of ≤ 25 decibels (dB) hearing level (HL) at each octave frequency from 250 to 8000 Hz in both ears. Participants were recruited through the Undergraduate Research Participant Pool at York University and by word of mouth. They were provided with credit towards their first-year psychology course or were paid $15 per hour. Informed consent was obtained in accordance with the ethics review boards at York University and Baycrest and conformed to the standards of the Canadian Tri-Council Research Ethics guidelines.

#### Pre-test: Mnemonic Similarity Task (MST)

To ascertain participant’s visual behavioral discrimination abilities, we tested participants on the Mnemonic Similarity Task (MST) (12). It was administered on a laptop running Windows 7. Participants were randomly assigned one of two MST stimulus sets and then administered the task followed an established protocol (12, 18). During the study phase, participants viewed 128 color images of everyday objects for 2 s each, followed by a 0.5 s ISI. For each picture, participants indicated via button press whether the object depicted was primarily an outdoor item or indoor item. A test phase followed this study phase. In the test phase, participants were administered a surprise recognition memory test. They were randomly presented with 192 images, each onscreen for 2 s, followed by a 0.5 s ISI. The photographs at test included 64 targets (studied objects), 64 unrelated foils, and 64 similar lures. Participants had to classify via a button press whether the image was old, new, or similar to the items presented at study. Results showed that participants were able to identify 78.43% of the targets as old; 49.65% of the lures as “similar” of the time; and 78.81% of the foils as new. These results are comparable to the target/lure/foil accuracy rates of 81.2%/47.6%/82.0%, found in a group of 26 healthy adults 20-39 years of age tested by Stark et al. (18).

#### Stimuli

We used six auditory patterns (micropatterns) in the study. Each was comprised of a different temporal arrangement of five sounds (Figure 1). The micropatterns were structured using pure tones, 100 ms long, with a 10 ms fade in and 10 ms fade out to prevent click sounds associated with abrupt frequency changes. The tones had frequencies of 350 Hz, 500 Hz, 638 Hz, 720 Hz, and 920 Hz. The latter four frequencies were used by Schröger (1994) (13). All micropatterns were compiled using Audacity 2.3.3 (http://www.audacityteam.org). In pilot studies, we identified micropatterns that were both distinctive and similar enough for testing. For example, the standard micropattern’s contour was M-shaped; its tones rise in frequency, then drop, then rise again, then drop. In contrast, the deviant micropattern’s contour switched directions three times. There were three highly similar micropatterns, or lures, presented during the test phase. Lure one and lure two switched contour direction once, and lure three switched contour twice, but these changes were in the opposite direction of the standard. There was one easily distinguishable micropattern (the foil), first presented at test. The foil had a straight increasing contour from low to high. Its five tonal frequencies were pitched 12 semitones down, making it sound low and distinct from the other micropatterns.

#### Passive study phase: incidental memory encoding

We divided the experiment into two phases. During the initial, 25-minute study phase, the participants sat in a double-walled sound-attenuating booth while watching a movie with the volume muted. We instructed participants to pay attention to the movie. Our aim was to prevent attention-elicited ERP signals that might obscure the MMN. At the same time, participants were exposed to the auditory input of the standard and deviant micropatterns, presented in 700 and 300 trials, respectively. The standard and deviant micropatterns were separated by a jittered (900 – 1150 ms) interstimulus intervals (ISIs). During the study phase, ERPs for each presentation of a micropattern were recorded, and are described in more detail below.

#### EEG recording and analysis

We collected ERP data during the study phase of the experiment. The EEG was sampled using a 76-channel acquisition system (BioSemi Active Two, Amsterdam, The Netherlands) with a bandpass of 0.16-100 Hz and a sampling rate of 512 Hz. Electrode positions were based on the International 10–20 system. Horizontal and vertical eye positions were recorded by electrooculography using four electrodes positioned around each eye. Two additional electrodes were placed on the left and right mastoids. EEG recording was grounded by an active Common Mode Sense (CMS) electrode and a passive Driven Right Leg (DRL) electrode. EEG recordings were processed offline using Brain Electrical Source Analysis 7.0 software (BESA 7.0; MEGIS GmbH, Gräfelfing, Germany).

The EEG data were visually inspected to identify segments contaminated by defective electrode(s). Noisy electrodes were interpolated using values from the surrounding electrodes, and no more than eight electrodes were interpolated per participant. The EEG was then re-referenced to the average of all electrodes and digitally filtered with 1 Hz high-pass filter (forward, 6dB/octave) and 40 Hz low-pass filter (zero phase, 24 dB/octave). For each participant, a set of ocular movements was identified from the continuous EEG recording and then used to generate spatial components that best account for eye movements. The spatial topographies were then subtracted from the continuous EEG to correct for lateral and vertical eye movements as well as for eye-blinks. After correcting for eye movements, the EEG was then scanned for artifacts. The data were parsed into 500 ms epochs, including 100 ms of pre-stimulus activity, and those including deflections exceeding ± 60 µV were marked and excluded from further analysis. The remaining epochs were averaged according to electrode position and stimulus type. Each average was baseline-corrected with respect to a 200 ms pre-stimulus baseline interval. Approximately 5-10% of trials were rejected for each participant. The results from the time domain (i.e., ERPs) and distributed source analysis (see below) were exported into BESA Statistics 2.0 for statistical analyses. BESA Statistics 2.0 software automatically identifies clusters in time, frequency, and space using a series of *t*-tests that compared the ERP amplitude between experimental conditions at every time point. This preliminary step identified clusters both in time (adjacent time points) and space (adjacent electrodes) where the ERPs differed between the conditions. The channel diameter was set at 4 cm, which led to around four neighbors per channel. We used a cluster alpha of .05 for cluster building. A Monte-Carlo resampling technique (Maris & Oostenveld, 2007) was then used to identify those clusters that had higher values than 95% of all clusters derived by random permutation of the data. The number of permutations was set at 1,000.

The cluster-based statistics reveal significant difference between standard and deviant, with the two strongest MMN signals at ∼340 ms and ∼605 ms after the onset of the micropattern. Figure 2B shows the group mean ERPs elicited by the standard and deviant micropatterns, as well as the corresponding difference wave. For the purpose of correlation analyses, the standard and deviant amplitudes were calculated as a mean voltage at the 40 ms period centered at the peak latencies in the individual participant waveforms collected from six of the frontal electrodes (FCz, Fz, F1, F2, FC1, FC2), where the largest response was obtained. The MMN amplitude for each individual was calculated by subtracting the mean standard amplitude from the mean deviant amplitude.

#### Test phase: recognition memory test

The test phase assessed the recognition memory of the participants for the studied micropatterns. Participants were presented with a micropattern and asked, “Did you hear this tone during the movie?” Participants were then instructed to respond “yes” by pressing the left arrow key, or “no” by pressing the right arrow key. The six micropatterns (two old, three lures, and one foil) were presented ten times each over the course of the test phase, for a total of 60 trials, in randomized order, with a 500 ms interval between stimulus presentation and instruction screens. The correct response to hearing the standard or deviant micropatterns was the left arrow key. The correct response to the three lure micropatterns and the foil micropattern was the right arrow key.

#### Posttest: Same-different forced-choice discrimination task

We concluded our experiment with a discrimination task. Our intent was to ensure that the participants could discriminate the micropatterns from one another when they were presented back-to-back. During the task, two micropatterns were played in succession and participants were asked whether they were the same or different. Participants indicated their answers via an arrow key. Every micropattern was presented against itself and against every other micropattern. There were 42 unique combinations of different micropatterns (as the different pairs could be presented in different order). Trials were randomly presented twice, leading to 84 discrimination trials. One female participant was unable to complete this task; therefore, analysis was conducted on 30 participants.

Discrimination results from the discrimination phase are reported in Supplementary Table 1. Results are collapsed across 21 unique pairs, disregarding the order of presentation (e.g., Similar-Foil and Foil-Similar were treated as one unique pair). As can be seen, healthy young adults showed high discrimination rates (*M =* 96.63%, *SE* = 1.53%) in this post-test, with slightly higher overall rates for discriminating same stimuli as “same” over discriminating different stimuli as “different.”

#### Statistics

Recognition accuracy data, as well as comparisons between correct rejection and *d’* rates, were analyzed using paired *t*-tests (two-tailed or one-tailed *t*-tests as noted) in IBM SPSS Statistics Version 26. Effect sizes for the value of *t* were computed according to Rosenthal (19). When necessary, multiple comparison corrections were applied using the Holm-Bonferroni method (20). Sensitivity values (*d’*) were calculated according to Macmillan and Creelman (21). The relationship between *d’* sensitivity and the MMN1 amplitude was analyzed using bivariate Pearson correlations in SPSS. Confidence intervals for the correlation coefficients were computed using the bias corrected accelerated (BCa) option. As negative MMN values indicate a stronger mismatch signal, negative correlations reflect a positive, not a negative relationship between conditions of interest and the MMN amplitude.

#### ERP source analysis

BESA 7.0 was used to estimate distributed source activity for ERPs elicited by standard and deviant stimuli. To enhance the accuracy of our findings and to reduce noise, we used distributed source analysis for each participant and each condition. We modelled the standard and deviant events and then compared the strength of the source activity. We did so by using an iterative application of Low-Resolution Electromagnetic Tomography (LORETA), which reduces the source space in each iteration. This imaging approach, termed Classical LORETA Analysis Recursively Applied (CLARA), provides more focal localizations of the brain activity and can separate sources located in close vicinity. The voxel size in Talairach space was 7 mm; this default setting is appropriate for the distributed images in most situations. The regularization parameters that account for the noise in the data were set with a single value decomposition cutoff at 0.01%.

**Supplementary Table 1.** Discrimination accuracy (percent correct)

| Contrast | Correct response | Discrimination<br>accuracy (mean) | Discrimination<br>accuracy (SE) |
| --- | --- | --- | --- |
| Standard–Standard | Same | 96.67% | 1.55% |
| Deviant–Deviant | Same | 96.67% | 1.55% |
| Foil–Foil | Same | 95.83% | 1.70% |
| Lure1–Lure1 | Same | 99.17% | 0.82% |
| Lure2–Lure2 | Same | 98.33% | 1.64% |
| Lure3–Lure3 | Same | 99.17% | 0.82% |
| Average-Same |  | 97.64% | 1.35% |
| Standard–Deviant | Different | 97.50% | 1.81% |
| Standard–Foil | Different | 100.00% | 0.00% |
| Standard–Lure1 | Different | 87.50% | 4.04% |
| Standard–Lure2 | Different | 96.67% | 1.55% |
| Standard–Lure3 | Different | 83.33% | 3.97% |
| Deviant–Foil | Different | 97.50% | 1.81% |
| Deviant–Lure1 | Different | 94.17% | 2.55% |
| Deviant–Lure2 | Different | 99.17% | 0.82% |
| Deviant–Lure3 | Different | 95.83% | 2.07% |
| Foil–Lure1 | Different | 99.17% | 0.82% |
| Foil–Lure2 | Different | 100.00% | 0.00% |
| Foil–Lure3 | Different | 98.33% | 1.14% |
| Lure1–Lure2 | Different | 99.17% | 0.82% |
| Lure2–Lure3 | Different | 97.50% | 1.37% |
| Lure1–Lure3 | Different | 97.50% | 1.37% |
| Average-Different |  | 96.22% | 1.61% |
| Average-Both |  | 96.63% | 1.53% |

